# Separation of multiple motor memories through implicit and explicit processes

**DOI:** 10.1101/2021.05.27.445962

**Authors:** Gefen Dawidowicz, Yuval Shaine, Firas Mawase

## Abstract

Acquisition of multiple motor skills without interference is a remarkable ability in sport and daily life. During adaptation to opposing perturbations, a common paradigm to study this ability, each perturbation can be successfully learned when a dynamical contextual cue, such as a follow-through movement, is associated with the direction of the perturbation. It is still unclear, however, to what extent this context-dependent learning engages the cognitive strategy-based explicit process and the implicit process that occurs without conscious awareness. Here, we designed four reaching experiments to untangle the individual contributions of the explicit and implicit components while participants learned opposing visuomotor perturbations, with a second unperturbed follow-through movement that served as a contextual cue. In Exp. 1 we replicated previous adaptation results and showed that follow-through movements also allow learning for opposing visuomotor rotations. For one group of participants in Exp. 2 we isolated strategic explicit learning by inducing a 2-sec time delay between movement and end-point feedback, while for another group we isolated the implicit component using the task-irrelevant error-clamp paradigm, in which participants were firmly instructed to aim their reaches directly to the target. Our data showed that opposing perturbations could be fully learned by explicit strategies; but when strategy was restricted, distinct implicit processes contributed to learning. In Exp.3, we examined whether the learned motor behaviors are influenced by the disparity between the follow-through contexts. We found that the location of follow-through targets had little effect on total learning, yet it led to more instances in which participants failed to learn the task. In Exp. 4, we explored the generalization capability to untrained novel targets. Participants showed near-flat generalization of the implicit and explicit processes to adjacent targets. Overall, our results indicate that follow-through contextual cues influence activity of both implicit and explicit processes during separation of motor memories. Furthermore, the follow-through context might activate, in part, top-down cognitive factors that influence not only the dynamics of the explicit learning but also the implicit process.

## Introduction

Our extraordinary ability to learn multiple motor tasks without interference allows us to flexibly switch between different environments and maintain a broad motor repertoire. In ball sports for example, we can adjust the strength and direction of our throw based on environmental changes, such as the ball’s weight (e.g., volley ball vs. takraw ball), without drastically affecting our skill. Formation of a motor memory associated with any motor skill task is believed to transpire through small trial-by-trial corrections that eventually allow learning to accumulate. This learning process is comprised of multiple distinct components (Huberdeau et al., 2015; Mazzoni & Krakauer, 2006; Smith et al., 2006; Taylor et al., 2014), at least one of which is implicit, slow and sensitive to sensory prediction error (McDougle et al., 2015; Ryan Morehead et al., 2017; Wolpert & Flanagan, 2001), and another process which is explicit, fast and sensitive to target error (Heuer & Hegele, 2011; Krakauer et al., 2000; McDougle et al., 2015; Morehead & Orban De Xivry, 2021; Poh & Taylor, 2019; Taylor et al., 2014; Werner & Bock, 2007).

Although great advances have been made in previous work in attempt to understand the contributions of the implicit and explicit processes (Taylor et al., 2014) and their interaction (Miyamoto et al., 2020) in motor learning, most of these works focused on learning of a single perturbation (Kim et al., 2018; Miyamoto et al., 2020; Taylor et al., 2014). Whether a similar parallel conclusion can be drawn regarding the contributions of the implicit and explicit processes when simultaneously learning multiple perturbations is still unclear. The generalization of the implicit/explicit dissection to more complex movements is essential to better understand motor behavior in real world conditions (Poh et al., 2021; Sarwary et al., 2015; Schween et al., 2018).

A common paradigm to study the ability to simultaneously learn multiple tasks is sensorimotor adaptation to opposing perturbations, such as two opposing force field perturbations. In this extreme scenario, when direction of perturbation switches randomly from trial to trial (Karniel & Mussa-Ivaldi, 2002; Howard et al., 2015;; Sheahan et al., 2016), the opposing learning directions interfere and neither perturbation is learned. This interference, however, can be markedly reduced when appropriate contextual cues, like associating each perturbation with a unique subsequent follow-through movement (Howard et al., 2015; Sheahan et al., 2016), can segregate learning of the opposing perturbation into distinct motor memories.

Here, we sought to explore whether dynamic follow-through contextual cues allow separation of motor memories through explicit processes, implicit processes or both. We designed reaching experiments and manipulated the implicit and explicit components while participants learned opposing visuomotor perturbations (clockwise and counterclockwise) that were randomly selected for each trial, with a second unperturbed follow-through movement. We isolated the implicit component by introducing error-clamp trials, a method that has been proven to successfully eliminate development of explicit strategies during visuomotor adaptation (Ryan Morehead et al., 2017). To isolate the explicit process, we used the 2-sec cursor endpoint feedback delay paradigm. This technique has been shown to minimize the use of the implicit component during visuomotor adaptation (Brudner et al., 2016; HELD et al., 1966; Schween et al., 2014; Schween & Hegele, 2017), in part by delaying the input from the cerebellar olivier nucleus to the cerebellar cortex (Ekerot & Kano, 1989; Herzfeld et al., 2018). Next, we examined whether the learned motor behaviors of implicit and explicit components are influenced by the disparity of the follow-through contexts. To do so, we decreased the distance between the follow-through movements associated with each perturbation and tested its effect on adaptation. Finally, we tested the generalization of learned movements to novel untrained directions. We hypothesized that follow-through contextual cues during adaptation to opposing visuomotor rotations prevent interference and that both the explicit and implicit processes contributed to overall learning. When contextual follow-through cues partially overlap, the ability to separate opposing memories will be reduced. Finally, generalization of the implicit process will be local and centered around the implicit plan whereas generalization of the explicit process is uniform across the novel directions.

## Methods

### Subjects

Ninety-one young right-handed healthy people, informed about the tasks involved in the experiments, but naïve to the objectives of the study, with normal or corrected-to-normal vision were recruited to the study (52 females, 39 males, age=26±4.13 years). Exclusion criteria included any neurological problems, motor dysfunctions, cognitive dysfunctions, uncorrected visual impairments, and/or orthopedic problems that would hinder reaching movements and/or affect the ability of the subject to understand and perform study tasks. Participants were recruited from the student population of the Technion – Israel Institute of Technology, Haifa, Israel, and gave written informed consent to participate in the study, which was approved by the local ethical committee of the Technion.

### Apparatus and general experimental procedures

Participants were seated approximately 60 cm in front of a reaching task system setup which consisted of a digitizing tablet and stylus pen (62 × 46 cm; Intuos4, Wacom Co., Japan) and a computer screen (48 cm width, 1280 × 1024-pixel resolution) which was reflected onto a semi-mirror that was positioned horizontally in front of the subject in order to obscure the view of the subject’s own hand and forearm during reaching tasks (Fig. 1A). The 2-dimensional position of the hand was continuously recorded by the tablet at a rate of 144 Hz. Participants made fast reaching movements from one of two possible square starting positions (3×3 mm), first toward a central circular target (2 mm diameter) and then toward a follow-through circular target (2mm diameter) which appeared at ±45° in experiments 1 & 2 & 4 (see Figures 1, 2 and 4 respectively), or ±10° in experiments 3 & 4 (see Figures 3 and 4 respectively). The distance between the starting position and the central target, as well as between the central target and follow-through target, was 10 cm. Auditory feedback based on movement time (MT) of the movement from the starting position to the central target was given as a high-frequency tone for movements that were too fast (MT<175 ms) and a low-frequency tone for movements that were too slow (MT>575 ms). If the movement was within the desired time frame (175≤MT≤575), no audio feedback was played. After each trial, the starting position appeared again and the participants returned to it, however, the cursor representing their hand movements would only appear when the participants were within 2 cm of the starting position.

**Figure 1.**
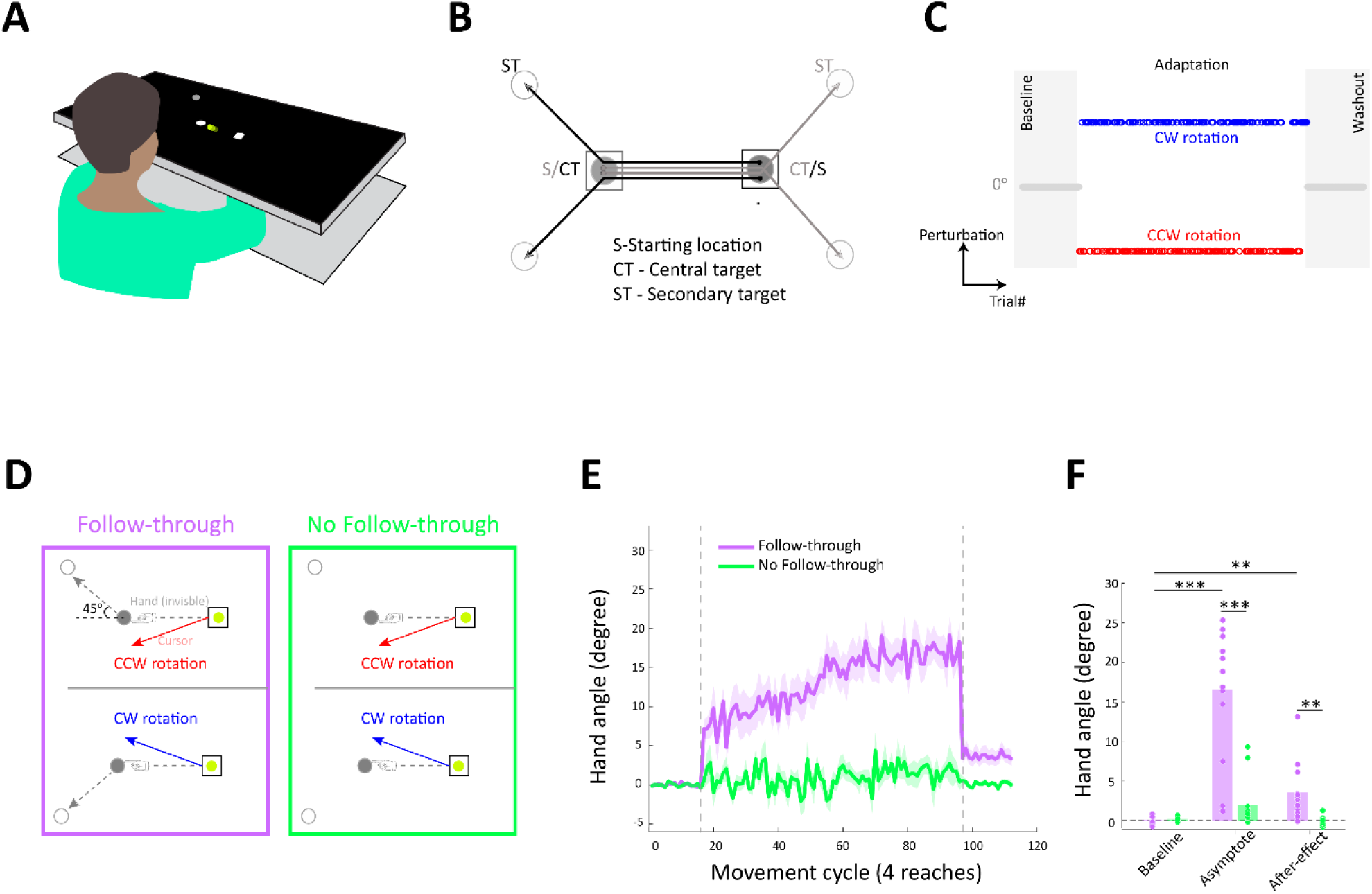
Experimental setup, protocol and finding of Experiment 1. **A.** Illustration of the experimental setup. Participants were seated in front of a reaching task system setup which consisted of a digitizing tablet and stylus pen and a computer screen which was reflected onto a semi-mirror that was positioned horizontally in front of the subject in order to obscure the view of the subject’s own hand and forearm. **B.** A schematic view of hand trajectories (during baseline). Color indicates the location of the second target relative to the movement to the central target; red 45º clockwise (CW) and blue counter-clockwise (CCW). **C.** The experiment consists of three stages: Baseline, Adaptation and Washout. The direction of the rotation was in the opposite direction of the second target and rotation sign changed randomly from trial to trial. **D.** Schematic representation of task structure in both groups of Experiment 1. Participants made initial movement to a central target (gray circle). While both targets were visible to both groups (gray and white circles), only the follow through group continued the movement to the second target. On exposure trials, visuomotor rotation (solid arrow) was applied during the initial movement. The direction of the rotation was applied in the opposite direction of the secondary target. **E.** Mean hand trajectory angle (the sign of the responses to the CW rotation was flipped) across subjects of each group, *follow-through* and *no follow-through*. Shading denotes SEM. **F.** Bars indicate mean hand trajectory angle in each block: baseline, asymptote and after-effect. Dots are individuals.

**Figure 2.**
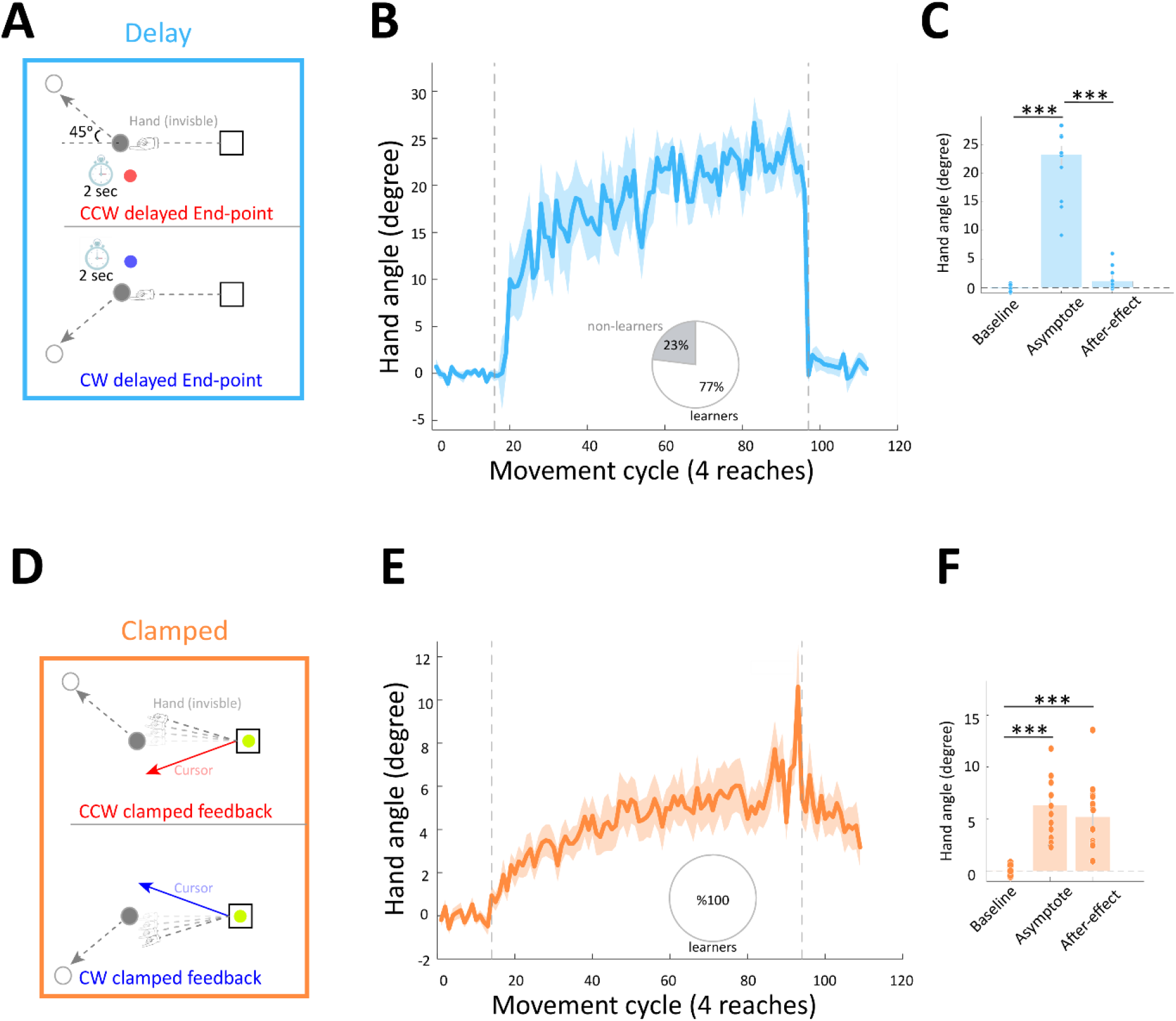
Isolating explicit and implicit components during learning opposing motor memories. **A.** Schematic representation of task structure for the delayed group of Exp. 2. The delayed group performed the full follow-through task, but we removed the online feedback of the cursor and instead provided it as an end-point after a delay of 2 sec. **B.** Mean hand trajectory angle across subjects in the delayed group. Inset shows the percentages of learners and non-learners. **C.** Bars indicate mean hand trajectory angle of the delayed group in each block: baseline, asymptote and after effect. Dots are individuals. **D.** Schematic representation of task structure for the clamped group of Exp. 2. Participants in that group were introduced to a task-irrelevant error-clamp visual feedback and instructed to continue aiming for the central target and to ignore the cursor manipulation. **E** and **F.** Similar to B and C but for the clamped group.

**Figure 3.**
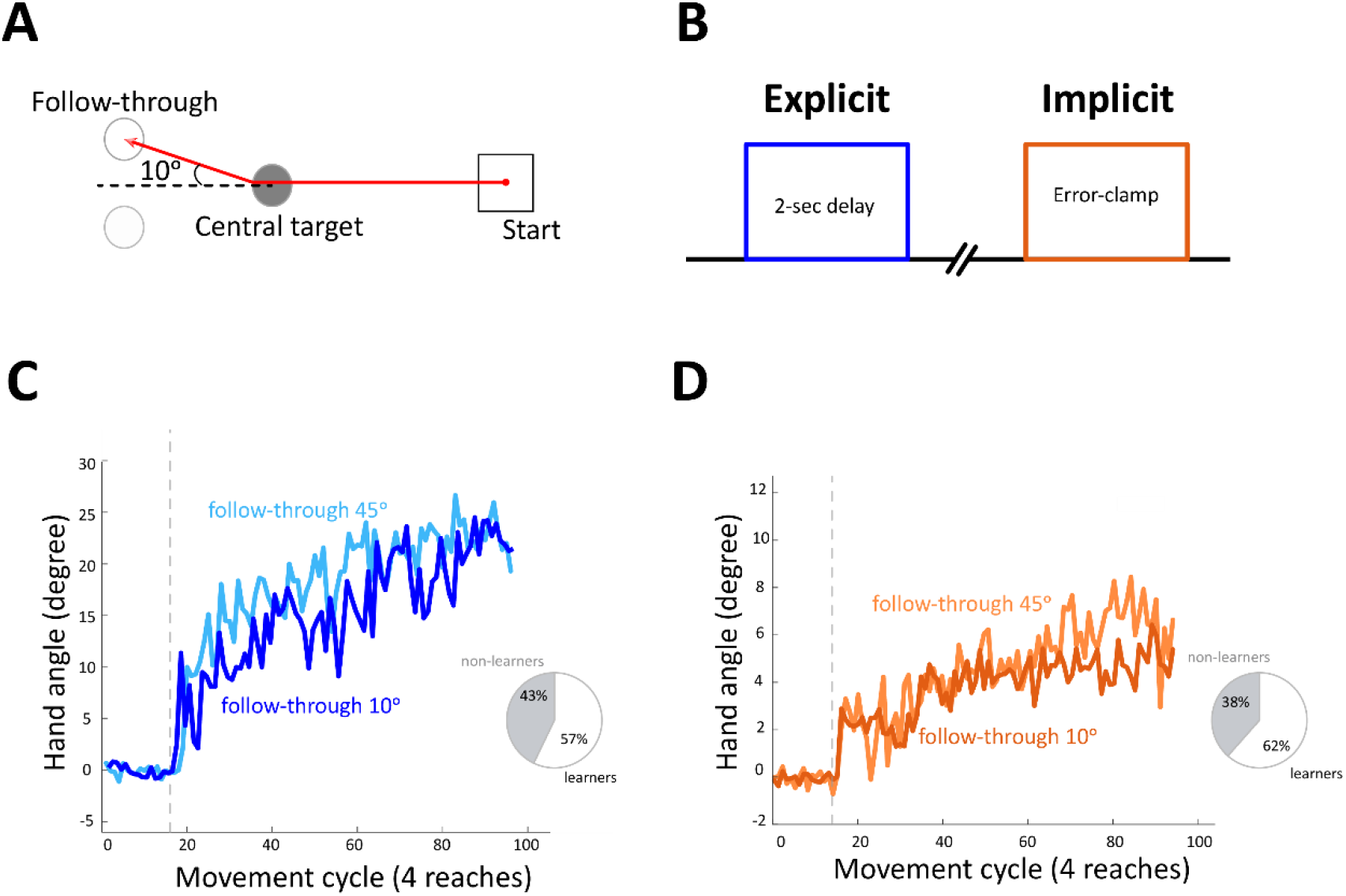
The influence of spatial distance of follow-through cues on separation of motor memories. **A.** Schematic representation of task structure for Exp 3. Secondary targets were moved closer together to ±10° from central target. **B.** Exp 3 protocol; participants performed explicit (2-sec delay) and implicit (error-clamp) blocks. **C.** Mean hand angle across subjects of explicit section of Exp. 3 (*follow-through 10°*) compared to mean hand trajectory angle across subjects of delayed group of Exp 2. (*follow-through 45°)* with chart of learners and non-learners of Exp.3. **D.** Mean hand trajectory angle across subjects of implicit section of Exp. 3 (*follow-through 10°)* compared to mean hand trajectory angle across subjects of clamped group of Exp. 2 (*follow-through 45°)* with chart of learners and non-learners.

**Figure 4.**
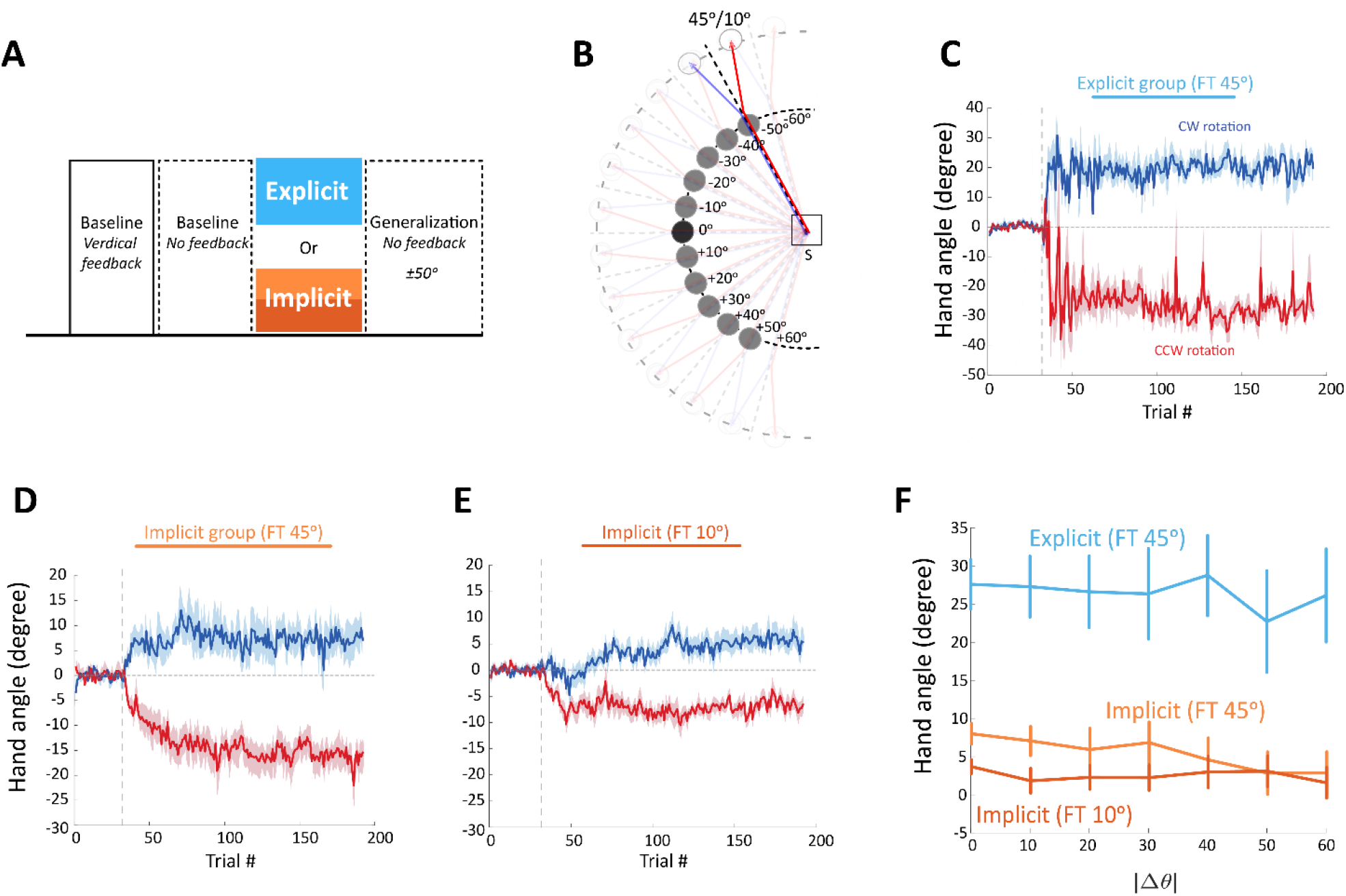
Explicit and implicit generalization of follow-through context. **A.** Exp. 4 protocol. Participants performed two baseline blocks (one with feedback and one without) followed by either the explicit experiment of Exp. 3 or the implicit experiment of Exp. 3. The last block was a generalization block with no feedback. **B.** Schematic representation of task structure of the generalization (and baseline) block of Exp.4. **C.**Mean hand trajectory angle across subjects of explicit group of Exp.4, movements with CW rotation in blue and movements with CCW rotation in red. **D.** Similar to C but for implicit group with follow-through targets at ±45°. **E.** Similar to C and D for implicit group with follow through targets at ±10°. **F.** Generalization functions. Hand angle of all groups as a function of the distance from learned target (|Δθ|).

### Experiment 1 – The effect of follow-through movements on formation of visuomotor memories (n= 25)

Experiment 1 was designed to examine the effect of subsequent follow-through movements in the ability to learn randomly-switched opposing visuomotor rotations. The cohort for Exp.1 was split up into 2 groups, a *follow-through* group (n=13, 7 women, mean age 26±4.91 years, range 20-35) and a *no follow-through* group (n=12, 9 women, mean age 25±3.16 years, range 22-31). The *follow-through* group was requested to make full follow-through movements toward the follow-through targets immediately after reaching the central target, while the *no follow-through* group received a visual cue of a follow-through target but were instructed not to move toward it. Thus, in this group, the participants only executed the movement to the central target. Both groups in this experiment performed three blocks of trials: *Baseline*, *Adaptation* and *Washout*. The *Baseline* block included 64 trials with no perturbation. Participants were then allowed a short rest period (1-2 minutes) before starting the next block. The *Adaptation* block included 320 trials and consisted of the same targets as *Baseline*, but the visual feedback presented on the screen was different. In this block, during the first movement (e.g., toward the central target), there was a ±30° visuomotor rotation between cursor and hand movement, randomly switched between + and - across trials. The sign of the rotation in each trial specified whether a clockwise or counter-clockwise (CW or CCW, respectively) rotation was applied. In order to reach the target, the participant had to adjust his/her hand trajectory to correct for the perturbation. Crucially, each perturbation was associated with the appearance of a single follow-through target. For example, reaching toward the left central target under a +30° rotation was always associated with a follow-through target that appeared at −45° to the midline, while reaching under a - 30° rotation was always associated with a follow-through target that appeared at +45° to the midline (Fig. 1D).

Participants in both groups were explicitly instructed that the aim of the experiment is to hit the central target with the cursor. Central and follow-through targets appeared at the same time on the screen and before participants initiated their movements. In both groups, visual feedback of the cursor was only given throughout the first movement, but not for the follow-through movement. All participants then received another break (4-5 minutes) before proceeding to the final block. In the *Washout* block (64 trials), the cursor feedback was entirely removed and the participants were instructed to continue aiming for the target as they did in the previous blocks. Again here, participants from the *follow-through* group were requested to make full follow-through movements toward the follow-through targets immediately after reaching the central target, while participants in the *no follow-through* group were instructed not to move toward it.

### Experiment 2 – The contribution of explicit and implicit processes in separation of motor memories (n= 21)

Experiment 2 was designed to isolate the effects of the explicit and implicit processes in separation of motor memories. To do so, one group of participants (*clamped group*, n=8, 5 women, mean age 30±6.34 years) was introduced to a task-irrelevant error-clamp visual feedback and instructed to continue aiming for the central target and to ignore the cursor manipulation. That is, while moving to the central target, the cursor showed a fixed trajectory of ±30° from the central target (+30° fixed rotation for a follow-through target at −45° to the midline, and −30° fixed rotation for a follow-through target at +45°), a manipulation which limits the strategic explicit component (i.e., strategy-free) to better isolate the implicit component (Kim et al., 2018; Ryan Morehead et al., 2017). To isolate the effect of the explicit process during learning, a second group of participants (*delayed group*, n=13, 8 women, mean age 26±1.97 years) performed the full follow-through task but the online feedback of the cursor was removed and instead provided as an end-point after a delay of 2 sec, a manipulation which limits implicit motor adaptation to better isolate strategic-based explicit learning (Brudner et al., 2016; McDougle & Taylor, 2019). Location of central and follow-through targets were identical to Experiment 1 and participants in both groups were instructed to make full follow-through movements toward the follow-through targets immediately after the movements toward the central target.

### Experiment 3 – The influence of spatial distance of follow-through cues on separation of motor memories (n= 16)

We examined whether the learned motor behaviors of the explicit and implicit processes are influenced by the disparity between the follow-through contexts. To test this, the spatial locations of the follow-through targets were set closer to each other at ±10 degrees from the central target (instead of ±45 as in Experiments 1 & 2). Here, all participants (n=16, 8 women, average age 26±6.04 years, range 18-38) performed one session of implicit error-clamped trials followed by an explicit delayed-feedback session. Adaptation sessions were separated with no-perturbation trials (128 trial washout session) in order to verify that any residual learned behavior decayed back to baseline level. The order of the sessions was counterbalanced across participants such that half of the participants started with the explicit delayed-feedback session while the other half started with the implicit error-clamp session.

### Experiment 4 – Explicit and implicit generalization of follow-through context (n=29)

Here we aimed to assess the generalization of the explicit and implicit learning to novel untrained directions. Participants were pseudo-randomly assigned either to *explicit* or *implicit* training condition groups. Each condition began with three no-perturbation Baseline blocks. The first block consisted of movements toward 13 different probe targets (0°, ±10°, ±20°, ±30°, ±40°, ±50° and ±60°), followed by subsequent follow-through targets at ±45° from each of the central target’s midline. Participants were provided with online cursor feedback for the first movement only but still instructed to make movements toward the follow-through target. The second Baseline block was identical to the first, but the visual feedback was removed. This was done to assess baseline directional biases at the different probe locations with no feedback. Then participants performed the third Baseline session, which included 64 trials, but only toward the central target located at 0° with visual feedback for the first movement. In the subsequent adaptation phase (320 trials), a ±30° visuomotor rotation was introduced to a single training movement direction located at 0°. Participants in the *explicit* condition group (n=8, 4 women, average age 25±3.33 years, range 21-30 years) received 2-sec delayed end-point feedback of their movements, whereas participants in the *implicit* condition group (n=11, 6 women, average age 25±3.29 years, range 19-26 years) received a fixed ±30° error-clamp feedback. In both conditions, the sign of the rotation was determined based on the location of the follow-through targets, as done in previous experiments. After the Adaptation block, participants performed a short Washout block (4 trials) to assess after-effects. Here, the cursor was removed in both groups and participants were instructed to aim for the central target. Next, participants in the *implicit* condition group performed a short Re-adaptation block (8 trials) in order to reattain the asymptotic level of the learned behavior that may have decayed during the Washout block. Lastly, participants performed a Generalization block (52 trials) which consisted of movements toward the 13 probe targets, followed by subsequent follow-through targets at ±45° from each of the central targets. Participants in the *explicit* condition performed three rounds of the short Re-adaptation block (6 trials), followed by the Generalization block (26 trials), in order to remain at the asymptotic level of the learned behavior as the explicit learning decays faster. No visual feedback about the cursor was provided in the Generalization blocks for either group. An additional control group (n=10, 8 women, average age 25±4.12 years, range 21-33) performed the same blocks as the *implicit* group, but the subsequent follow-through targets were located at ±10° from each of the probe central targets.

### Data analysis

Kinematic data was collected from the tablet at 144 Hz and stored on a computer for off-line analysis using MATLAB (The MathWorks, Natick, MA). Movement performance was quantified at each trial using *hand trajectory angle* (°), defined as the angle between the imaginary lines connecting the movement origin to the movement completion location and the movement origin to the target location. Positive values indicated clockwise angles whereas negative values indicated counterclockwise angles. For each participant, the hand trajectory angle was sign-adjusted appropriately so that errors from CW and CCW rotation trials could be combined and then binned in epochs of 4 consecutive movements. Performance at different phases of the adaptation curve were calculated in order to quantify overall learning. First, learning curves were normalized by subtracting the average hand trajectory angles of the four baseline epochs just prior to rotation onset to account for any incomplete washout. Then, “*Baseline*” performance was defined as the mean hand trajectory angle of the last four epochs in the Baseline block, the “*Asymptotic*” level of learning was defined as the mean hand trajectory angle of the last four epochs in the Adaptation block and the “*After-effect*” was defined as the mean hand trajectory angle of the first four epochs of the Washout block. The mean and standard error (SE) for each measure across participants was then calculated.

The amount of generalization (Exp. 4) was calculated by subtracting the mean hand trajectory angle of each direction in the second Baseline block from the mean hand trajectory angle across trials in the last Generalization block of that direction, to correct for any directional biases that might exist when removing the visual feedback of the cursor in theses blocks. Here also, for each participant, the hand trajectory angle was first sign-adjusted appropriately so that angles from +45°/+10° follow-through and −45°/−10° follow-though trials could be combined, respectively.

### Statistical analysis

To perform the statistical comparison and analysis in Exp. 1, we performed a 2-way ANOVA repeated measure with a main factor of experimental group (follow-through vs. no follow-through) and learning epochs (Baseline, Asymptotic, After-effect) as the second level. Post-hoc comparison between groups at different epochs was conducted using a two-tailed t-test. Post-hoc comparison between epochs within groups was conducted using a two-tailed paired t-test. In Exp. 2, we used a separate 1-way repeated measure ANOVA comparing the different learning epochs for the implicit and explicit conditions, as we were interested in examining the isolated effect of each component of the total learning. In Exp. 3, we used a repeated measure 2-way ANOVA with a main factor of experimental condition (explicit vs. implicit) and learning epochs as the second level. We used repeated measure in this experiment because all participants performed the two conditions. In Exp. 4, for each group of participants, we first averaged the generalization pattern across the follow-through directions (i.e., +45° and −45°, +10° and −10°, for group 1,2 and 3, respectively) and then we ran a 1-way ANOVA with a main factor of probe targets (0°, ±10°, ±20°, ±30°, ±40°, ±50° and ±60°). The generalization level at each direction was tested with single-sample t-tests, with the test variable set at 0. Significance level for all tests was set at 0.05.

#### Learners vs. Non-learners

Based on previous reports using similar paradigms (Brudner et al., 2016) and pilot data from our lab, we expected to find some participants that might exhibit minimal changes in their behavior in response to the rotation and therefore fail to learn the tasks. To objectively distinguish between participants who learned and those who did not, we computed a one-sample *t-test* on individual participant data sets, comparing the hand trajectory angle in the last four epochs (*Baseline*) of the Baseline block and the last four epochs of the rotation session (*Asymptotic*). A participant that reached a significant difference (*p* < 0.01) for this measure was defined as a learner, otherwise she/he was classified as a non-learner. Of note, we presented only the results (e.g., hand trajectory angle, group-based analysis) of the learners.

## Results

### Experiment 1

In this experiment, one group of participants (*follow-through group, n=13)* made a subsequent unperturbed movement to a secondary follow-through target, whereas a second group (*no follow-through group, n=12)* only made reaching movements toward the central target. Follow-through movement was critical for separation of opposing memories of visuomotor rotations. Statistically, there was a significant group effect on learning epochs (repeated measures two-way ANOVA: F(1,22)=37.917, p<0.0001, *η*^2^ = 0.17671, Fig. 1F) and a significant learning epochs × group interaction (F(2,44)=21.959, p<0.0001, *η*^2^ = 0.1907). The change in hand trajectory angle from the Baseline (0.0595±0.4936°) to the end of the Adaptation block (i.e., asymptote, 16.5455±8.1998°) was significantly larger (paired t test: t_12_=7.3573, p=8.76*10^−6^, Cohen’s d=2.838) for the *follow-through* group but not for the *no follow-through* group (paired t test: t_10_=1.4956, p=0.1656, d=0.629). These differences were also evident in the after-effect measure, taken from the first 4 epochs of the no-feedback Washout block (see Materials and methods). When comparing after-effects vs. baseline epochs, the *follow-through* group showed significant (paired t test: t_12_=3.3098, p=0.0062, d=1.39) after-effects (3.5739±3.537°) whereas no after-effect (0.0032±0.5606) was observed in the *no follow-through* group (paired t test: t_10_=−0.7726, p=0.4576, d=0.314). These results replicated previous studies in force-field adaptation and showed that contextual cues in the form of follow-through movement allow learning of visuomotor skills that otherwise interfere.

### Experiment 2

Data of Experiment 1 cannot disambiguate the relevant contribution of explicit and implicit processes to the net learning. The fact that the *follow-through* group showed significant, yet small, after-effects and the clear discrepancy between total learning and after-effects, suggests that follow-through contextual cues allow learning opposing perturbations not only through strategy-based explicit processes. To better understand how explicit strategic and implicit learning interact during follow-through movement, we trained participants on a task that allowed us to isolate how each component evolves throughout the course of training.

Participants in the *delayed* group performed the full follow-through task but received feedback of the cursor as an end-point after a delay of 2 sec, a manipulation which limits implicit motor adaptation to better isolate strategic learning (Fig. 2A). Here, we found a rapid significant increase (paired t test: t_9_=10.0125, p=3.5411*10^−6^, d=4.4225) in the hand trajectory angle (asymptotic level of: 21.8852±7.0201°) during the rotations block, relative to baseline performance (−0.1433±0.5819°). Interestingly, when the rotation was abruptly removed, no significant after-effect (1.1082±2.4979°) was reported (paired t test: t_9_=1.5323, p=0.1598, d=0.6901). These results indicate that the *delayed* group learned to counter the opposing rotations over the course of the rotation block solely by developing explicit strategies with negligible contribution of the recalibration implicit processes as manifested by the absence of the after-effect.

The second group was introduced to a task-irrelevant error-clamp visual feedback (*clamped* group), a manipulation which limits explicit learning to better isolate implicit recalibration learning (Morehead et al., 2017). As depicted in Figure 2E, participants in this group showed a gradual but significant (paired t test: t_7_=3.8012, p=0.0067, d=1.9438) adaptation to the opposing rotations. Crucially, instruction to ignore cursor trajectory but make unperturbed follow-through movement enabled segregation of the memory into distinct implicit processes. Data showed increase in hand trajectory angle between baseline (−0.0968±0.284°) and asymptote (6.9019±5.084°) and significant (paired t test: t_7_=4.469, p=0.0029, d=2.3484) after-effect as reported early during the Washout block. Hand trajectory angle early in the Washout block (3.9023±2.3915°) was not different (paired t test: t_7_=2.2237, p=0.0615, d=0.755) from asymptote performance (6.9019±5.084°) (Figure 2F). Overall, this data suggests that when strategy is restricted, follow-through contextual cues allow participants, in large part, to separate opposing motor memories through implicit processes.

### Experiment 3

So far, our results have shown that spatially distinct follow-through movements allow separation of motor memories and that this segregation includes a large implicit component. Next, we addressed a follow-up question and examined whether directional distance between the follow-through cues affects the separation of the motor memories.

In Experiment 3, we reduced the spatial distance between the follow-through contexts (from ±45° to ±10° to the central target) and tested its effect on implicit and explicit components during learning opposing visuomotor rotations. Specifically, we examined whether the learned motor behaviors of each component, observed in previous experiments, is affected when the follow-through context is located at ±10° to midline from the central target. Here, participants performed a block of task-irrelevant error-clamp visual feedback trials and a block of 2-sec delay feedback trials (See Methods and Materials). In the delay block, participants showed rapid correction, compensating for the opposing visuomotor rotations. Asymptotic performance (18.4033±9.0537°) was significantly (paired t test: t_8_=5.9413, p=3.4532*10^−4^, d=2.9041) higher than late baseline performance (−0.2192±0.5186°). We noted, however, a small after-effect in this block (paired t test: t_8_=#x2212;2.4691, p=0.0388, Cohen’s d=−0.947) that seems to be driven by data from two participants and is not constant across all subjects. When we compare the performance during the Adaptation delay block between this group (follow-through at ±10°) and the delayed group from Experiment 2 (follow-through at ±45°), we found no clear differences. Asymptotic level (18.4033±9.0537° vs. 21.8852±7.0201°) and after-effect (1.198±1.3667° vs. 1.1082±2.4979°) were not significantly different (repeated measures two-way ANOVA: F(2,34)=0.556, p=0.577, *η*^2^ = 0.0044). Nevertheless, we found that reducing the spatial distance between the follow-through cues affected the number of learners. Here 29% of participants failed to learn the task whereas 23% from the delayed group in Experiment 2 failed to learn.

Next, we examined whether the implicit learning process is affected by the directional distance between the follow-through movements. We found that the majority of participants (62%) in the error-clamp block were able to implicitly learn the task by monotonically changing their hand movement even when the spatial distance between the follow-through targets is quite small. Asymptotic performance (6.2413±3.6711°) was significantly higher (paired t test: t_7_=5.0102, p=0.0015, d=2.4096) than late baseline performance (−0.0717±0.5014°). When we compare the performance during Adaptation between this group (follow-through at ±10°) and the error-clamp group from Experiment 2 (follow-through at ±45°), we found no significant differences. Asymptotic level (6.2413±3.6711° vs. 6.9019±5.084°) was not significantly different (repeated measures two-way ANOVA: F(1,14)=0.940, p=0.761, *η*^2^ = 0.0015) between experiments. However, we found again that reducing the spatial distance between the follow-through cues affected the number of learners. Here, 38% of participants failed to learn the task whereas none of the subjects from the delayed group in Experiment 2 failed to learn.

These findings suggest that the change in directional distance of the follow-through movements has a small effect on learning and/or performance levels of implicit and explicit processes, but it apparently affects the number of individuals that were able to learn the context-dependent perturbations.

### Experiment 4

Here, we tested the generalization pattern of explicit and implicit components by training participants with a single central target and two follow-through movements and examining movements to central targets at twelve other directions, but using similar follow-through contextual cues (i.e., ±45° to the midline of each central target). To test the generalization pattern of the explicit process, a group of participants (n=8) performed the 2-sec delay follow-through task in a single central direction and then tested in 12 new directions (6 CW and 6 CCW) (Fig. 4B). We found no significant effect of test target direction (1-way ANOVA: F(6,42)=0.577, p=0.746, *η*^2^ = 0.0054) on the percentage generalization. Generalization level on each direction was significantly larger than 0 (post-hoc t-tests: p<3.34*10^−6^ and d>2.154). Thus, an explicit adaptation to opposing perturbations learned with a single central-target direction leads to the acquisition of a transfer rule that generalizes uniformly across novel directions. These results indicate that the generalization of the explicit learning component is significantly modulated by a non-kinematic dimension of the follow-through context.

To test the generalization pattern of the implicit process, a new group of participants (n=11) performed the task-irrelevant error-clamp follow-through task in a single central direction and then tested in the 12 new directions (Fig. 4A and 4B). Interestingly, we found that there was no significant effect of test target direction (1-way ANOVA: F(6,54)=0.794, p=0.578, *η*^2^ = 0.0328) on the percentage generalization. In each direction, the generalization level was significantly larger than 0 (post-hoc t-tests: p<0.0112 and d>1.283). This result indicates that an implicit adaptation to opposing perturbations learned with a single central-target direction generalizes uniformly across novel directions with similar follow-through contextual cues. We note, however, that while small, there was a trend of negative slope in the implicit group with follow-through targets at +/−45°, suggesting that the implicit component might also be influenced by the movement-related features (i.e., movement direction). The finding of near-flat generalization pattern of the implicit process was replicated in an additional group (n=10) that implicitly learned to separate opposing motor memories but now with contextual follow-through cues at a closer distance of ±10° from the midline. Again, we found no significant effect of test target direction (1-way ANOVA: F(6,54)=1.034, p=0.413, *η*^2^ = 0.0446) on the generalization pattern, but in each direction, participants showed significant (post-hoc t-tests: p<0.015 and d>0.604) generalization (compared to null hypothesis of 0°). Overall, these results suggest that the generalization of the implicit learning component could also be modulated by a non-kinematic dimension of the follow-through context. This is a surprising finding because it is not in line with previous work that showed that the implicit learning component generalizes locally around the aim direction according to kinematic dimension (Brayanov et al., 2012; Krakauer et al., 2000; Poh et al., 2021; Poh & Taylor, 2019; Ryan Morehead et al., 2017).

## Discussion

Our experiments sought to isolate and understand the different components of adaptive learning during separation of opposing motor memories. In particular, we sought to dissociate the contribution of the implicit and explicit learning processes during learning randomly alternating opposing visuomotor rotations, each associated with a contextual follow-through movement. In the first experiment, we exhibited that it is difficult to learn opposing environments without dynamical contextual cues such as follow-through movements. In the second experiment, we found that strategic-based explicit processes explained most of the learned behavior during separation of motor memories. Yet, when strategic learning is restricted, the implicit process takes over and the opposing perturbations can be learned, but performance is saturated at low levels of learning. Furthermore, we found that reducing the distance between follow-through directions associated with each perturbation has little effect on total learning. Lastly, we separately explored the generalization function of explicit and implicit processes following learning with follow-through context and found near-flat uniform generalization across untrained directions of both components.

### Follow-through movements allow separation of opposing memories through explicit and implicit processes

The use of dynamic contextual cues such as follow-through or lead-in movements has been previously shown as a prerequisite condition which allows trial-by-trial separation of opposing motor memories in force-field adaptation (Howard et al., 2015). To understand whether this trial-to-trial separation process arises from compensation of the implicit component, strategic-based explicit component or both learning components, we performed multiple experiments while participants adapted to opposing visuomotor rotations. We used the visuomotor paradigm because it systematically dissociates the relevant contribution of each components and its response to external perturbations (Huberdeau et al., 2015; Mazzoni & Krakauer, 2006; Smith et al., 2006; Taylor et al., 2014). Here, we reported that participants could fully learn to compensate for the opposing rotations, when a follow-through contextual cue is available, entirely by developing an explicit strategy. The level of the asymptotic performance as well as the absence of an after-effect in the delayed group in Experiment 2 supports the idea that participants explicitly utilized the follow-through cues to separate memories that otherwise interfere. This is consistent with previous work that showed that during adaptation to opposing rotations, each associated with a distinct visual workspace cue, participants near fully compensated for the perturbation by developing a strategy, as probed by verbal reports (Taylor et al., 2014). Although in previous work explicit strategy was estimated using verbal aiming reports, we believe that our 2-sec delay protocol engaged similar strategic-based explicit mechanisms that are sensitive to performance error (Brudner et al., 2016). In addition, our finding also corroborates previous suggestion based on neuroimaging study that separation of distinct perturbations with different contexts might rely on cognitive components (Imamizu et al., 2003).

We also found evidence for trial-by-trial implicit learning during the separation process. Our data proposed that a follow-through context-dependent separation process is not exclusively driven by the explicit process, but it also engaged an implicit component that responds to the follow-through contextual cues. The presence of an after-effect of the full follow-through group already hints to the involvement of an implicit component during the separation process. In this block, no visual feedback was provided and participants were instructed to stop using any strategy they might have developed during the adaptation period. At this stage, it was impossible to determine whether this implicit process was compensating for sensory-prediction error (i.e., the difference between an action’s outcome and an internal prediction of the outcome), performance error (i.e., the difference between action outcome and the task goal) or both errors (Lee et al., 2018; Mazzoni & Krakauer, 2006; Taylor & Ivry, 2011), since all of these forms of error coexisted in the full follow-through group. The fact that we saw implicit learning when the strategic explicit process was constrained (no performance error) during the task irrelevant error-clamp condition, and similar magnitude of after-effects observed in this group compared to the full follow-through group, therefore indicates that implicit learning in our experiments is driven by sensory-prediction errors. Comparable after-effects however, do not necessarily imply similar underlying implicit learning. Our results cannot confirm if the same implicit process observed in the full follow-through group also played a role in the task-irrelevant error-clamp group. In addition, we confirmed a previous finding of incomplete learning of the implicit process as depicted in all groups of the task-irrelevant error-clamp condition. This lower asymptote of the implicit component is consistent with previous reports (Kim et al., 2018), suggesting that this phenomenon probably reflects a balance between learning from errors and forgetting of the adaptive state from one trial to the next (Shmuelof et al., 2012; van der Kooij et al., 2015)

Recent reports suggested that the implicit process could also be driven, at least in part, by performance errors and responds to strategic explicit processes (Albert et al., 2020). This is in line with previous studies that showed that strategy use interferes with the build-up of implicit adaptation (Benson et al., 2011; Jakobson & Goodale, 1989), as strategy use would decrease the performance errors that could drive implicit adaptation. For example, recent work has elegantly dissociated the interaction between the implicit and explicit learning from their responses to the external perturbation and demonstrated that implicit adaptation effectively compensates for noise in explicit strategy. This interaction raises the question of whether similar behavior as reported here can be drawn when both implicit and explicit processes are simultaneously co-active during the separation process. More recent findings proposed that implicit and explicit processes are not independent as previously thought, but rather coexist and interact during motor adaptation tasks (Mazzoni & Krakauer, 2006; Miyamoto et al., 2020). Future works is needed to explore the interaction between the implicit and explicit learning when both are simultaneously engaged during learning opposing environments.

### How do explicit vs. implicit learning processes relate to context-dependent separation of motor memories?

One way to think of the explicit learning process, in relation to adaptation to opposing rotations using follow-through context, is that it reflects deliberative caching of stimulus-response contingencies (McDougle & Taylor, 2019). That is, participants learn a specific stimulus (e.g., a CW follow-through movement in our experiment) each associated with a single rotation, then they map this stimulus into a distinct response, compensating for the perturbation. This discrete stimulus-response contingency reflects a type of strategic process that appears to be related to working memory. Evidence suggests that performance in a spatial working memory task correlates with the use of explicit strategies in visuomotor rotation learning (Christou et al., 2016; Seidler et al., 2012). In addition, the performance on a spatial working memory test correlated with the rate of early visuomotor learning, and both recruited a similar neural network (Anguera et al., 2010). The stimulus-response contingency in this abstract fashion, however, cannot fully explain our data, in particular, the absence of improved performance in the no follow-through group in Experiment 1. In this experimental condition, the stimulus was statically illustrated but no actual follow-through movement occurred and participants failed to deliberately associate this cue with the sign of the perturbation. Thus, the inability of the stimulus-response contingency to explain the behavior observed in our experiments, suggests that an additional process(es) sensitive to dynamic contextual cues must be involved in the learning process.

But why are contextual cues that require some movement elements crucial for separation of motor memories? Recent study by Sheahan and colleagues (Sheahan et al., 2016) demonstrated that planning a distinct follow-through movement is more important than execution in allowing separate motor memory formation. That is, information about the follow-through movement must be available during planning and before the initial movement is executed in order to dissociate the motor memories and facilitate learning of opposing force-fields. The importance of motor planning in learning challenging environments that often interfere was already reported by Hirashima and Nozaki (Hirashima & Nozaki, 2012), when they showed that opposing force-field motor memories can be learned and flexibly retrieved, even for physically identical movements, when distinct motor plans in a visual space were linked to each field. Altogether, previous work and ours suggest that separation of motor memories appears to depend not only on explicit contextual cues, but also on whether these cues engage actual planning of movement associated with the cue. This plan-based learning theory can fit adequately with a recent neural framework of a dynamical systems perspective of motor cortex (Ames et al., 2014; Kaufman et al., 2014). Within this framework, it seems likely that distinct planned follow-through movements bring the motor cortical population activity to two distinct initial preparatory states, which lead into two separate dynamical trajectories during movement. This hypothetical explanation about the link between plan-based learning and initial states of preparatory space of the dynamical neural system remains, however, unresolved.

### Contextual follow-through generalization of implicit and explicit processes

Generalization is a fundamental aspect of sensorimotor learning as it allows flexible transfer of what has been learned from one context to another. Here we tested the generalization pattern of explicit and implicit learning processes during learning opposing visuomotor rotations, each linked to a follow-through movement that served as a contextual cue. Our finding of near-flat explicit generalization is in line with recent work that showed that explicit learning is likely to produce relatively global generalization. For example, Heuer and Hegele (Heuer & Hegele, 2011) showed that participants reported similar rotated aims to adjacent targets, suggesting that their explicit estimation of the movement required to counteract the perturbation generalizes globally. Furthermore, Bond and Taylor (Bond & Taylor, 2015) showed that explicit learning is highly flexible and that participants have a more abstract representation of the aiming solution rather than just remembering the appropriate aiming landmark, again supporting the theory of global generalization of explicit processes. As discussed above, one way to think of explicit generalization, in relation to adaptation to opposing rotations using a follow-through context, is that it reflects generalization of deliberative caching of stimulus-response contingencies (McDougle & Taylor, 2019). In our study, we propose that the follow-through contextual cues during the delay condition might have engaged top-down inference about which action participants ought to take in a given follow-through context. That is, participants generalized strategies they had developed in the learned direction to a novel direction using information stemming from the follow-through contextual cues. A very recent study showed that part of the typical motor generalization function can be driven by distances between contexts in the psychological space (e.g., shape of the target) (Poh et al., 2021).

The implicit generalization in our experiments, however, provided surprising results. Although the magnitude of generalization in our implicit groups was smaller than the generalization of the explicit group, we reported that participants showed a near-flat global component of generalization to adjacent targets. This finding cannot be fully explained by the theory of plan-based local generalization of implicit processes. This plan-based theory is supported by recent results that found that the plan, not the movement itself, is the center of generalization (Day et al. 2016, McDougle et al., 2017). That is, participants showed local generalization that peaked around where they reported their aim, not around the task goal or movement direction, during the adaptation phase. If this was the case in our task-irrelevant error clamp groups, we should have been seen dramatic reduction of generalization on central targets located at ≥ ± 40°. Our results did not support this theory. Instead, it seems that the presence of the contextual follow-through cue in the novel direction affected, to some extent, the generalization pattern of the implicit process. One possibility that might explain this finding is that the implicit process is influenced not only by low-level kinematic features like movement direction, but also by some abstract psychological feature that was inferred by the follow-through contextual cues. The fact that participants in the implicit groups made follow-through movements and uniformly generalized in the novel directions, suggested that follow-through context provided some abstract psychological information that directly affected the implicit process.

In summary, our data proposed that follow-through contextual cues might not purely reflect traditional movement representation sensitive to directional distance between the cues. Instead, in our perspective follow-though context could represents other dimensions in movement space or even a mixture with high-level cognitive representation. Indeed, recent work by Poh and colleagues (Poh et al., 2021), showed that motor generalization in visuomotor adaptation tasks, is influenced by a mixture of at least two factors, kinematically-linked implicit representations (e.g., direction of target) and cognitive non-kinematic top-down inference (e.g., shape of target). It is possible that the improving performance in untrained directions during implicit learning is caused in part, by effect of cognitive non-kinematic top-down inference.

## Acknowledgements

This study was funded by the Israel Science Foundation (grant # 1634/19), and the German-Israel Foundation (grant # I-2535-409.10/2019).

